# Comprehensive Identification of Pathogenic Microbes and Antimicrobial Resistance Genes in Food Products Using Nanopore Sequencing-Based Metagenomics

**DOI:** 10.1101/2023.10.15.562131

**Authors:** Annie Wing-Tung Lee, Iain Chi-Fung Ng, Evelyn Yin-Kwan Wong, Ivan Tak-Fai Wong, Rebecca Po-Po Sze, Kit-Yu Chan, Tsz-Yan So, Zhipeng Zhang, Sharon Ka-Yee Fung, Sally Choi-Ying Wong, Wing-Yin Tam, Hiu-Yin Lao, Lam-Kwong Lee, Jake Siu-Lun Leung, Chloe Toi-Mei Chan, Timothy Ting-Leung Ng, Franklin Wang-Ngai Chow, Polly Hang-Mei Leung, Gilman Kit-Hang Siu

## Abstract

Foodborne pathogens, particularly antimicrobial-resistant (AMR) bacteria, remain a significant threat to global health. Conventional culture-based approaches for detecting infectious agents are limited in scope and time-consuming. Metagenomic sequencing of food products offers a rapid and comprehensive approach to detect pathogenic microbes, including AMR bacteria. In this study, we used nanopore-based metagenomic sequencing to detect pathogenic microbes and antimicrobial resistance genes (ARGs) in 260 food products, including raw meat, sashimi, and ready-to-eat (RTE) vegetables. We identified *Clostridium botulinum* and *Staphylococcus aureus* as the predominant foodborne pathogens in the food samples, particularly prevalent in fresh, peeled, and minced foods. Importantly, RTE-vegetables, which harbored *Acinetobacter baumannii* and *Toxoplasma gondii* as the dominant foodborne pathogens, displayed the highest abundance of carbapenem resistance genes among the different food types. Exclusive *bla^CTX-M^*gene-carrying plasmids were found in both RTE-vegetables and sashimi. Additionally, we assessed the impact of host DNA and sequencing depth on microbial profiling and ARG detection, highlighting the preference for nanopore sequencing over Illumina for ARG detection. A lower sequencing depth of around 25,000 is adequate for effectively profiling bacteria in food samples, whereas a higher sequencing depth of approximately 700,000 is required to detect ARGs. Our workflow provides insights into the development of food safety monitoring tools and can assess the potential risk to human health from foodborne pathogens and ARGs. This approach has the potential to revolutionize the screening of food products and enable more efficient and accurate detection of foodborne pathogens and ARGs, thereby reducing the risks of foodborne illness and improving public health.

## Introduction

Antimicrobial resistance (AMR) is a major threat to public health worldwide in the 21^st^ century (Prestinaci, Pezzotti et al. 2015). While most research has focused on the burden of AMR in clinical settings, recent statistical reports show that infections acquired in the community outnumber those acquired in hospitals (Jin, Xie et al. 2022). Agricultural settings, where high doses of antibiotics are often prescribed to prevent and treat infections in food animals, have been identified as hotspots for AMR. The transmission of foodborne pathogens, particularly AMR foodborne bacteria, is facilitated by animals and plants, which serve as primary reservoirs and mediators (Founou, Founou et al. 2016). This poses a significant global threat to public health, as foodborne illnesses, such as Salmonellosis, Campylobacteriosis, and Vibrio infections, may not respond to antibiotic treatment, leading to severe symptoms such as diarrhea, dehydration, and even death (Solomon and Oliver 2014, Mendelson and Matsoso 2015, Fao 2016).

To control the risks of foodborne illness, microbial profiling of food products is an effective method for detecting infectious agents in food. However, conventional culture-based approaches, are time-consuming and limited in scope, only detecting less than 1% of the bacterial diversity (Pedron, Guyon et al. 2020, Sala-Comorera, Caudet-Segarra et al. 2020, Marshall, Kurs-Lasky et al. 2021). Recently, several sequencing techniques have been employed to improve food safety, including metabarcoding, shotgun sequencing (in which DNA fragments of 300 to 500 base pairs are used), and long-read sequencing (Billington, Kingsbury et al. 2022). Metabarcoding, which is lower in cost, requires prior knowledge of the microbial and antibiotic resistance gene (ARG) targets for targeted sequencing (Billington, Kingsbury et al. 2022). This typically involves applying only one approach, such as 16S rRNA, ITS, 26S rRNA, or ARGs in each preparation. In contrast, long-read sequencing has a higher error rate than short-read technologies but has a simpler and more cost-effective workflow (Kovac, Bakker et al. 2017). It is worth noting that these sequencing techniques cannot reliably differentiate between living and dead organisms. However, the detection of nonviable cells can still indicate past contamination events that can trigger source-tracking investigations (Billington, Kingsbury et al. 2022). Additionally, these techniques can detect viable bacteria that cannot be cultured and microbes capable of producing toxins such as fungi (Bruno, Sandionigi et al. 2019, Koutsoumanis, Allende et al. 2019, Billington, Kingsbury et al. 2022).

Nanopore sequencing has emerged as a powerful long-read sequencing tool for microbial evaluation in various fields, including environmental and clinical applications (Schmidt, Mwaigwisya et al. 2016, Martin, Stebbins et al. 2021, Urban, Holzer et al. 2021, Chavan, Sarangdhar et al. 2022). This platform can generate reads up to 2 × 10^6^ bases in length, allowing for the accurate identification of ARGs and bacterial species with a 92-94% accuracy rate in mock communities (Benitez-Paez, Portune et al. 2016, Wang, Huang et al. 2020, Lao, Ng et al. 2022). Despite persisting concerns about the accuracy of nanopore sequencing at the base level, the technology has already made a significant impact on real-time tracing and rapid data sharing during outbreaks of bacterial and viral pathogens (Quick, Loman et al. 2016, Faria, Kraemer et al. 2018, Boykin, Sseruwagi et al. 2019, Urban, Holzer et al. 2021). Recent studies have attempted cost-effective and timely metagenomic-based investigations of foodborne outbreaks using bacterial spike-in food samples (Buytaers, Saltykova et al. 2021, Maguire, Kase et al. 2021). This resource is valuable for the surveillance of multidrug-resistant bacteria and the implementation of infection control in the food industry, which can help prevent the spread of AMR and foodborne infectious diseases.

In this study, we conducted a comprehensive assessment of the workflow, identifying foodborne pathogens and ARGs in different types of food and food processing methods. We also investigated the impact of sequencing depth, host DNA, and sequencing platform (Nanopore vs. Illumina sequencing) on the microbiome and AMR profiling results. This study provides valuable insights for the development of rapid risk analysis tools to assess the potential risk to human health from foodborne AMR microorganisms.

## Results

### Experimental workflows

The study employed nanopore-based metagenomic sequencing to analyze five different types of food, namely beef, chicken, pork, sashimi and ready-to-eat (RTE) vegetables. In brief, the food samples were initially homogenized using a stomacher and then centrifuged to separate the microbes from the food. This process took approximately two hours (Figure 1). Subsequently, DNA was extracted from the microbial samples using the QIAamp BiOstic Bacteremia DNA Kit, which took an additional 45 minutes. The entire sample preparation process for the nanopore sequencing, including the use of the Rapid Barcoding Kit (SQK-RBK004) in library preparation, took approximately four hours in the laboratory, with an additional 12 hours required for nanopore sequencing (Figure 1).

**Figure 1.**
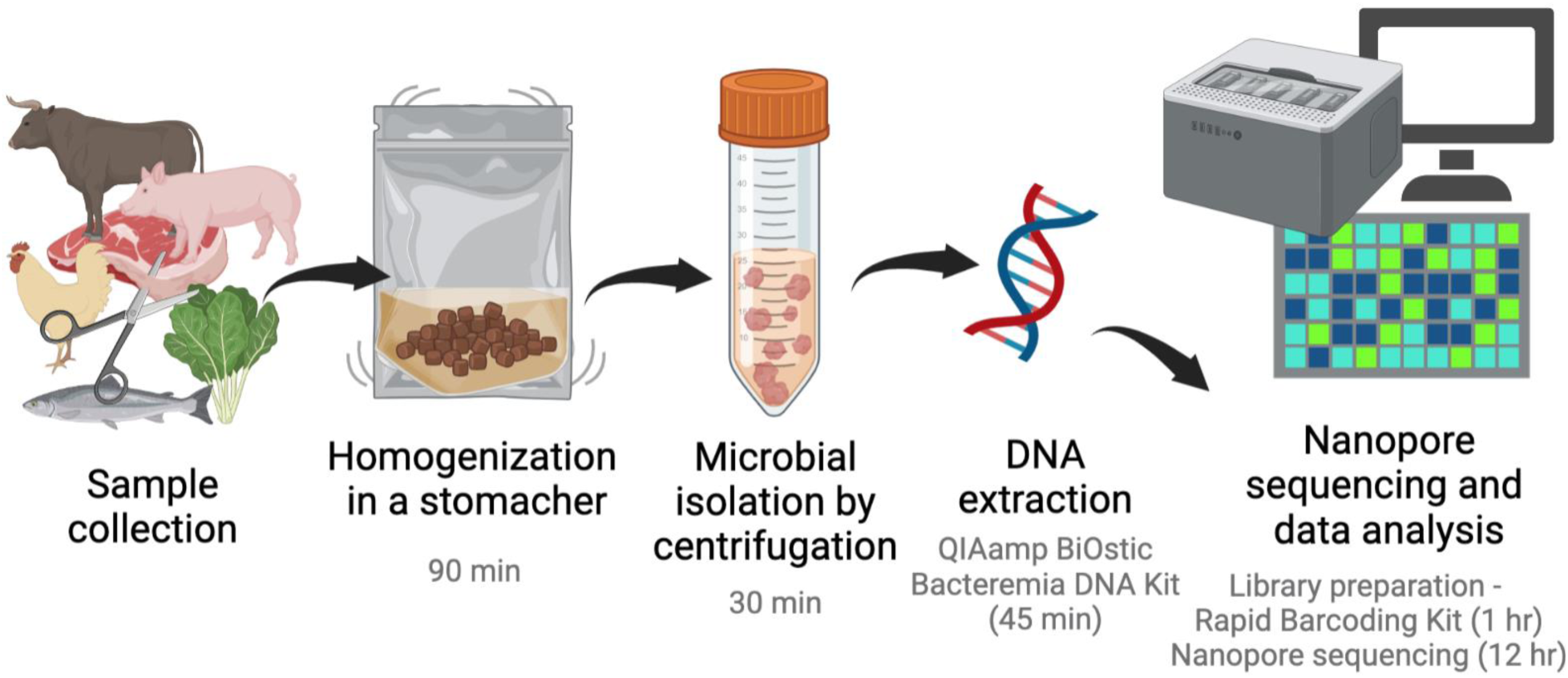
Food microbiome study design and workflow. For nanopore-based metagenomic sequencing, five different types of food were sampled, including beef, chicken, pork, sashimi, and ready-to-eat (RTE) vegetables. The food samples were initially homogenized using a stomacher, followed by centrifugation to separate the microbes from the food, which took around two hours. Subsequently, DNA was extracted from the microbial samples using the QIAamp BiOstic Bacteremia DNA Kit, which took an additional 45 minutes. The entire sequencing process on the nanopore platform, including the use of the Rapid Barcoding Kit (SQK-RBK004), took approximately four hours in the laboratory, with an additional 12 hours dedicated to nanopore sequencing. The diagram was generated using Biorender.

### Overall microbial community in food

A total of 137,404 ± 129,391 reads (>1,000 bp) were obtained from each sample (Figure 2A). The metagenomic sequencing successfully identified the taxonomic compositions of bacterial, archaeal, fungal, protozoal, and viral microorganisms in 260 food products. The analysis revealed that bacteria (61.5% ± 8.1%) was the most abundant microbial group in the five food types, followed by protozoa (16.5% ± 10.6%), fungi (8.2% ± 3.7%), viral (8.2% ± 2.2%), and archaea (5.5% ± 1.6%) (Figure 2B, Supplementary Data 1). When comparing the abundance of microbial groups in different food types, it was observed that archaea had the highest prevalence in RTE-vegetables, while bacteria were most abundant in chicken. Fungi and viruses were found to be dominant in sashimi, whereas protozoa were the dominant group in pork (Figure 2B).

**Figure 2.**
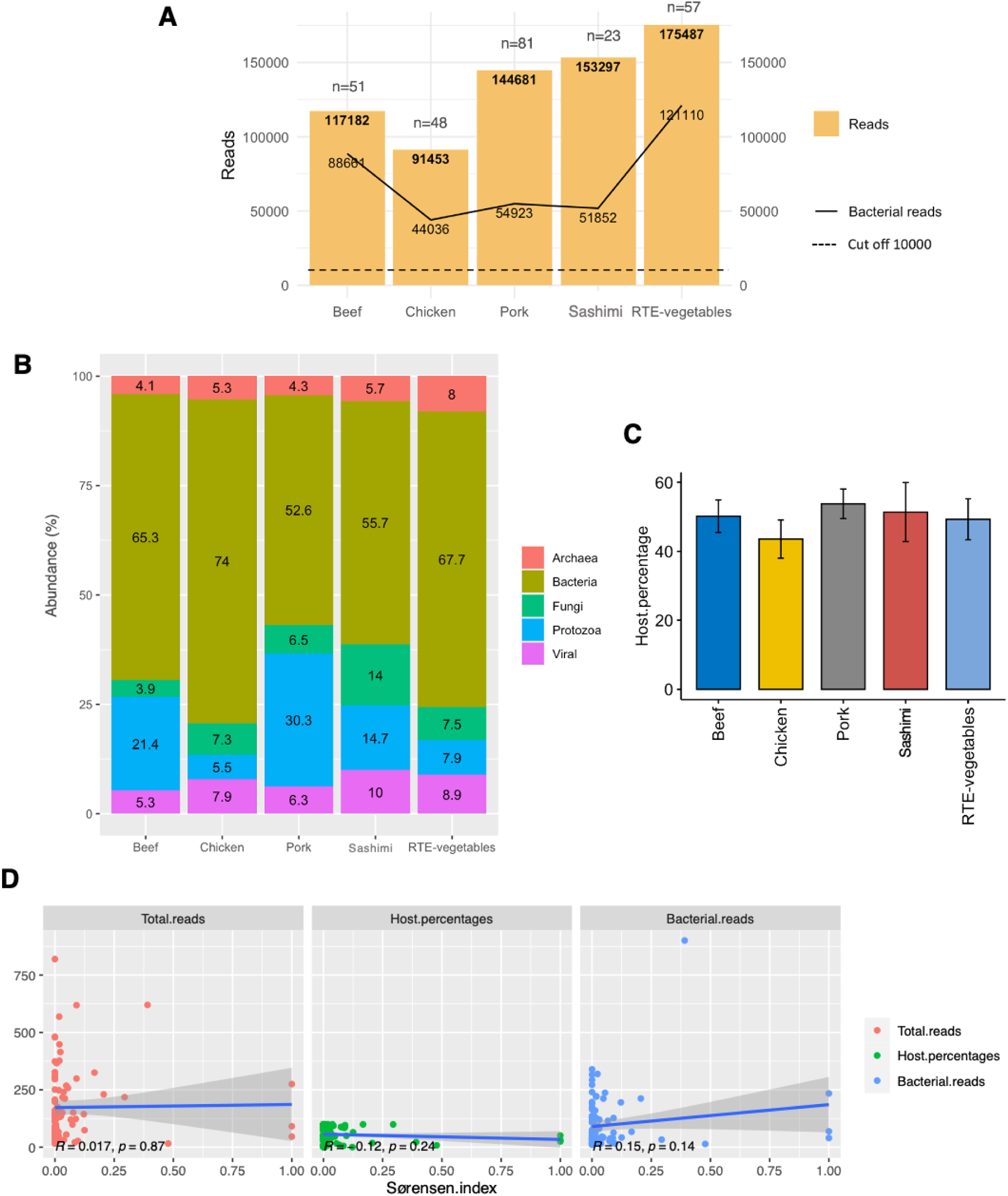
Microbial communities in beef (n=51), chicken (n=48), pork (n=81), sashimi (n=23) and RTE-vegetables (n=57). (**A**) Read depth and bacterial classification summary. The bar plot shows the total number of reads and the number of bacterial reads. The dashed line indicates the cut-off at 10,000 reads. (**B**) Taxonomic compositions of the five food products include bacteria, archaea, fungi, protozoa, and viruses. (**C**) The mean percentage of host DNA found in each food product. Data presented as mean ± SEM. (**D**) The Sørensen–Dice index was calculated for the bacterial dataset (species of relative abundance of <1% were discarded) with and without host DNA and the relationships between the Sørensen–Dice index and total reads (per 1000 reads), host percentages (%) and bacterial reads (per 1000 reads) were shown.

Figure 2C presents the average proportion of host DNA detected in each food sample (n=102, 50.2% ± 37.6%). The similarity of microbiota between datasets with and without host DNA was examined by computing the Sørensen–Dice coefficient (Figure 2D). The findings indicated that removing host DNA had a minimal impact, with Sørensen–Dice coefficients of 0.06 ± 0.18 (Figure 2D). No association was observed between the Sørensen–Dice index and total reads, host percentages and bacterial reads (Figure 2D, R-value < 0.2).

### Bacterial communities in the five food types

The principal component analysis (PCA) in Figure 3A demonstrated that the bacterial communities in the five food types were distinct (Supplementary Data 1), with RTE-vegetables showing the most significant difference. The correlation microbiome analysis also indicated that RTE-vegetables had a weak correlation (Figure 3B, coefficient value < 0.5) with all other food types. Notably, beef showed a weak correlation with chicken, sashimi, and RTE-vegetables (Figure 3B, coefficient value < 0.5), but a strong correlation with pork. In contrast, sashimi and chicken exhibited a strong correlation (coefficient value > 0.5), which was also observed in the PCA (Figure 3A and 3B).

**Figure 3.**
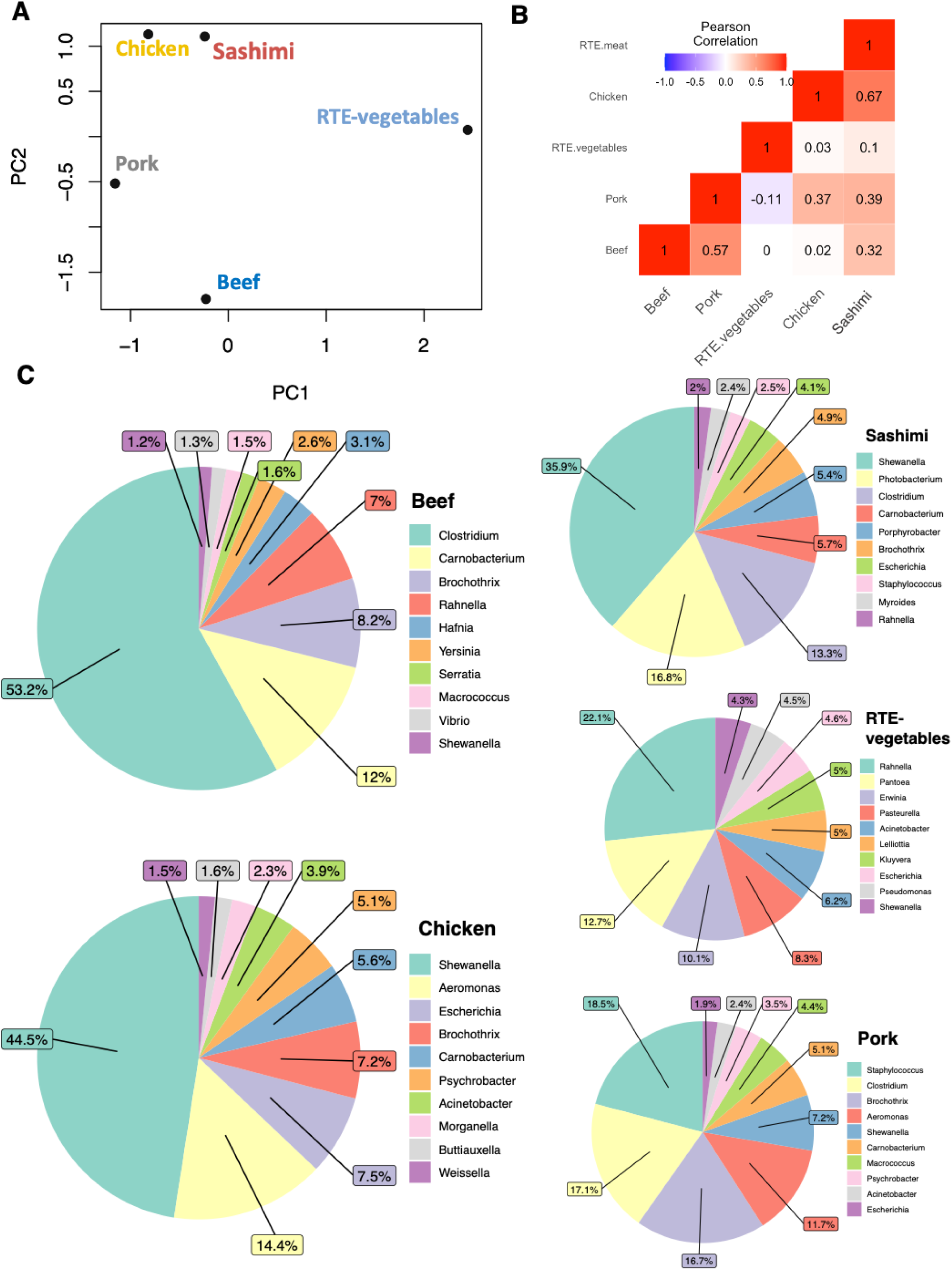
Microbial compositions of different food products. (**A**) Principal component analysis of the bacterial community in the five food samples at species level. Each dot represents one food type. (**B**) Correlation bacterial microbiomes analysis among food products. (**C**) The top 10 most abundant genera in food products were shown in the pie charts.

The dominant bacterial genera varied across the different food types. In beef, *Clostridium* (53.2%), *Carnobacterium* (12.0%), and *Brochothrix* (8.2%) were the most prevalent genera (Figure 3C). For chicken, the most abundant genera were *Shewanella* (44.5%), *Aeromonas* (14.4%), and *Escherichia* (7.5%). Sashimi was dominated by *Shewanella* (35.9%), followed by *Photobacterium* (16.8%) and *Clostridium* (13.3%) (Figure 3C). In pork, *Staphylococcus* (18.5%), *Clostridium* (17.1%), and *Brochothrix* (16.7%) were the most abundant genera (Figure 3C). In RTE-vegetables, *Rahnella* (22.1%), *Pantoea* (12.7%), and *Erwinia* (10.1%) were the most prevalent genera (Figure 3C).

### Signature species for each food type

A multi-level pattern analysis was carried out, employing the indicator value index to pinpoint distinctive species associated with each food type (Figure 4A, Supplementary Data 2). These distinctive species exhibited a statistically significant variation in their abundance (*p < 0.05*) and possessed an indicator value index exceeding 0.25. Hierarchical clustering of the signature species revealed a core microbiome in respective food products (Figure 4A). To illustrate, in beef, we identified *Clostridium botulinum* and *Carnobacterium divergens* as the distinctive species, while *Staphylococcus aureus* predominated in pork (Figure 4A). Chicken exhibited numerous distinctive species, including *Shewanella sp., Aeromonas sp.,* and *Escherichia coli* (Figure 4A). Sashimi was found to host *Photobacterium damselae* and *Porphyrobacter sp.* as its distinctive species, while RTE-vegetables displayed several distinctive species, such as *Pseudomonas fluorescens*, *Pasteurella multocida*, and *Pseudomonas azotoformans* (Figure 4A).

**Figure 4.**
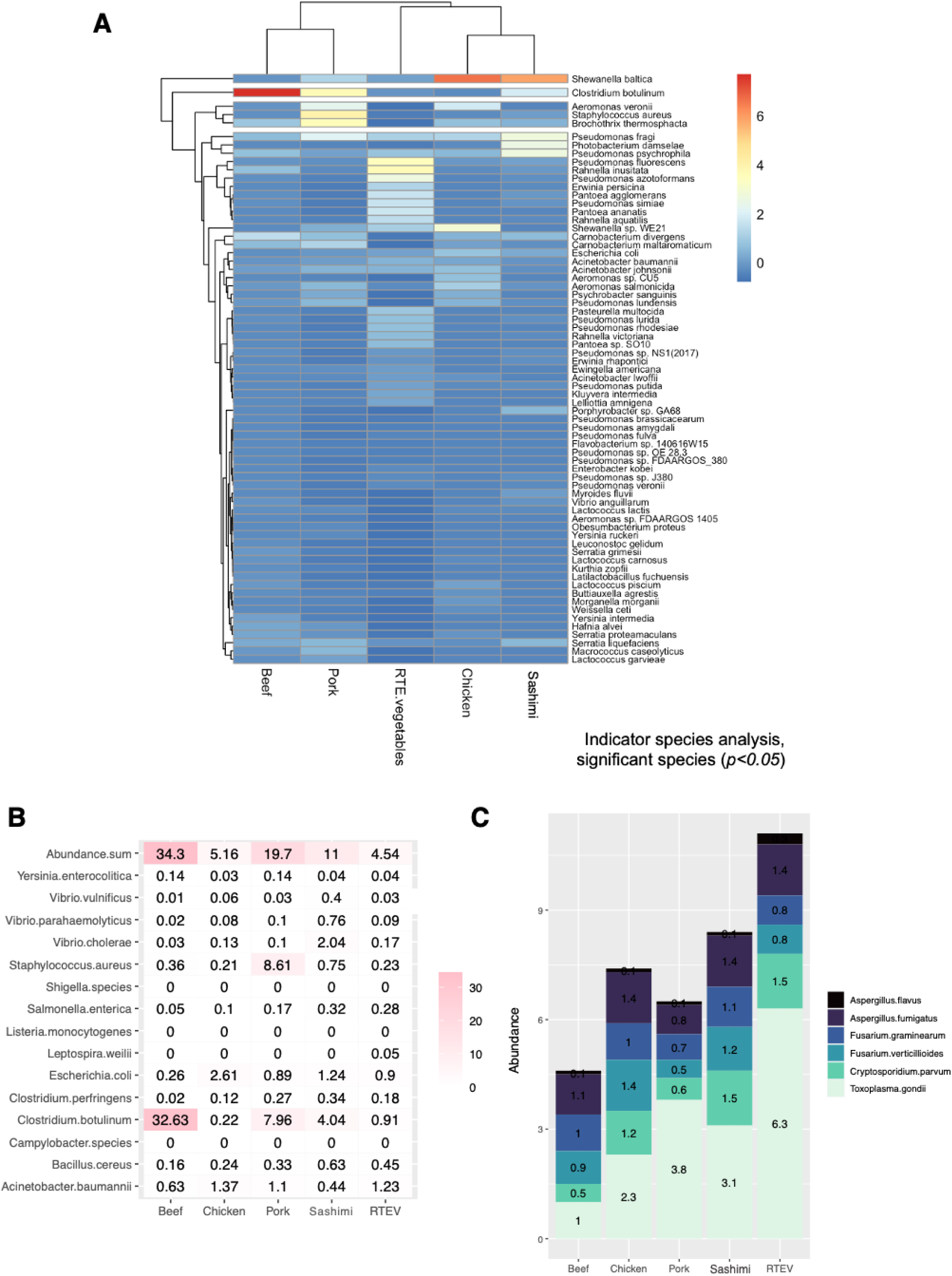
Signature and pathogenic for each food type. (**A**) Signature species that serve as distinctive indicators for each food type. (**B**) The mean abundance of 15 potentially pathogenic bacteria in various food products. The scale bar shows the mean abundance of the pathogenic bacteria (%). (**C**) Abundance of *Aspergillus, Fusarium, Toxoplasma gondii* and *Cryptosporidium parvum* in different food types.

### Identification of bacterial pathogens in different food types

We identified 15 common foodborne pathogens in the food products. In total, 85.4% (222/260) of the food samples contained pathogen reads with a total relative abundance exceeding 1% (Supplementary Data 1). More than 95% of beef and pork samples and 87.5% of chicken samples harbored pathogen reads. For sashimi and RTE-vegetables, the percentage of samples with pathogen reads was 78.3% and 64.4%, respectively (Supplementary Data 1). In each food sample, the overall bacterial pathogen abundance was 15.6% ± 21.7% (Figure 4B). The average pathogen abundance was highest in beef (34.3%), followed by pork (19.7%), sashimi (11.0%), chicken (5.2%), and RTE-vegetables (4.5%) (Figure 4B).

Among these species, *Clostridium botulinum* (9.3%), *Staphylococcus aureus* (2.9%), *Escherichia coli* (1.1%), and *Acinetobacter baumannii* (1.0%) ranked as the top four across the food products, followed by *Bacillus cereus* (0.3%), *Vibrio cholerae* (0.3%), and *Clostridium perfringens* (0.2%) (Figure 4B). *Salmonella enterica* (0.2%), *Vibrio parahaemolyticus* (0.1%), *Yersinia enterocolitica* (0.1%), and *Vibrio vulnificus* (0.1%) were also identified. *Leptospira weilii* was detected in both pork and RTE-vegetables, albeit at an average abundance of less than 0.01% (Supplementary Data 1). Conversely, *Campylobacter species, Listeria monocytogenes*, and *Shigella species* were not found in any of the samples (Figure 4A). Notably, *Clostridium botulinum* showed the highest mean abundance in beef at 32.6%, followed by pork at 8.0%, and sashimi at 4.0% (Figure 4B). *Staphylococcus aureus* had its highest abundance in pork at 8.6%, while less than 1% was observed in other food types (Figure 4B). *Escherichia coli* exhibited a higher abundance in chicken at 2.6% and sashimi at 1.2%, but it was less than 1% in other food types. Moreover, *Acinetobacter baumannii* was found to be more than 1% abundant in chicken, RTE-vegetables, and pork (Figure 4B). Sashimi exhibited a distinctive pathogen pattern, with three *Vibrio species* being more abundant than other food types, particularly *Vibrio cholerae,* which accounted for 2% (Figure 4B).

In parallel, beyond bacteria, we uncovered the presence of other potential foodborne microorganisms, including protozoa like *Toxoplasma gondii* and *Cryptosporidium parvum*, as well as fungi such as *Fusarium verticillioides, Fusarium graminearum, Aspergillus fumigatus,* and *Aspergillus flavus*, all showing varying levels of abundance (Figure 4C and Supplementary Data 1). The overall abundance was highest in RTE-vegetables, followed by sashimi, chicken, pork, and beef (Figure 4C). Notably, *Toxoplasma gondii* exhibited a significantly higher abundance in RTE-vegetables at 6.3%. It was worth noting that the levels of *Aspergillus fumigatus, Fusarium verticillioides,* and *Cryptosporidium parvum* were markedly elevated in chicken when compared to beef and pork (Figure 4C).

### Detection of antimicrobial resistance genes in different food types

The number of antibiotic resistance genes (ARGs) detected in each sample is depicted in Figure 5A. In total, ARGs were present in 258 out of the 260 food samples (Supplementary Data 3). Among these, RTE-vegetables exhibited the highest quantity of ARGs, followed by pork, beef, chicken, and sashimi (Figure 5A). Because of the clear classification of ARG groups by ARGpore2, we used it to further investigate the prevalence of ARGs specific to various antibiotic classes across different food products, with a focus on genes associated with aminoglycosides, β-lactams, macrolides, quinolones, vancomycin, and tetracyclines (Figure 5B, Supplementary Data 4). Genes linked to tetracycline were most prevalent in pork, accounting for 45.3% of the total ARGs identified. Notably, chicken showed the highest abundance of β-lactam-related genes (10.9%), followed by pork (7.7%) and beef (4%) (Figure 5B). Similarly, chicken also exhibited the highest abundance of ARGs related to aminoglycosides (3.6%) and quinolones (2.5%) (Figure 5B). In the case of macrolides, pork had the highest ARG abundance (2.3%), followed by RTE-vegetables (1.6%) and chicken (1.0%) (Figure 5B).

**Figure 5.**
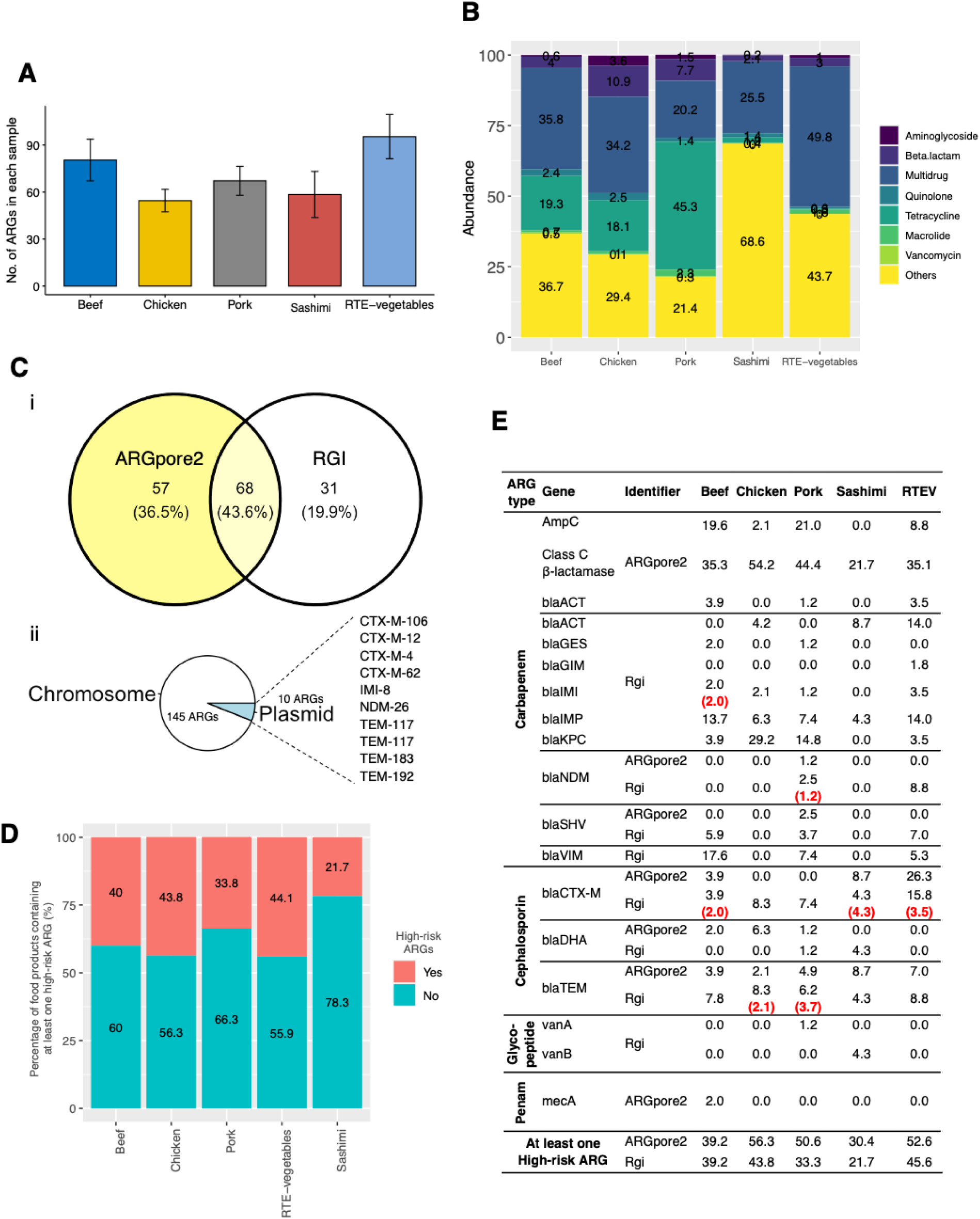
AMR composition in food products: beef (n=47), chicken (n=47), pork (n=76), sashimi (n=15) and RTE-vegetables (n=53). (**A**) Total number of ARGs detected in each sample. (**B**) Total abundance of genes related to aminoglycosides, macrolide, quinolone, tetracycline, beta-lactam, and multidrug in food products. (**C**) Detection of high-risk ARGs among the food samples. (i) Number of food samples carrying high-risk ARGs analyzed by ARGpore2 and RGI. (ii) Number of ARGs carried by plasmids and chromosome revealed by RGI. (**D**) Percentages of food products containing at least one high-risk ARG across various food types. (**E**) Percentages of food products containing the corresponding ARG. Red bracket represents the percentage of food samples carrying plasmid-mediated ARGs.

### Detection of high-risk antimicrobial resistance genes in different food types

In terms of high-risk ARGs, 60.0% (156 out of 260) of the food samples were found to carry at least one high-risk ARG, as indicated by the combined analysis with RGI and ARGpore2 (Figure 5Ci). Regarding the potential for ARGs to be transferred, only 10 were found to be linked with plasmids (Figure 5Cii). In each food type, the analysis by RGI revealed that 21.7% of sashimi samples and 44.1% of RTE-vegetable samples contained at least one high-risk ARG (Figure 5D). When examining raw meat samples, it was observed that the chicken group had the highest number of samples containing at least one high-risk ARG, at 43.8%, followed by beef at 40.0%, and pork at 33.8% (Figure 5D).

Meanwhile, Figure 5E provides a summary of the distribution of various types of ARGs in the food samples. We assessed high-risk ARGs, including carbapenem, cephalosporin, glycopeptide, and penam resistance genes, listing them alongside their respective detection methods, either ARGpore2 or RGI (Figure 5E). Specifically, according to RGI results, carbapenem resistance genes were most prevalent in RTE-vegetables at 57.9%, followed by beef at 45.1%, and chicken at 41.8% (Figure 5E). In the cephalosporin group, pork exhibited the highest abundance at 14.8%, followed by chicken at 11.7%. Additionally, regarding glycopeptide ARGs, *vanA* and *vanB* were detected in pork and sashimi, respectively, while the *mecA* gene was exclusively observed in beef (Figure 5E). Regarding plasmid-associated ARGs, as indicated by the red brackets in Figure 5E, most of them were *bla^TEM^ and bla^CTX-M^* genes. In particular, we found plasmid-associated *bla^CTX-M-4^, bla^CTX-M-12^,* and *bla^CTX-M-62^* in sashimi and RTE-vegetables (Figure 5D and E). Furthermore, all the identified *bla^IMP^* genes in beef samples were determined to be plasmid-mediated. Additionally, it was worth noting that 1.2% of pork samples contained plasmids carrying *bla^NDM^* (Figure 5E).

In comparing the results of ARGpore2 and RGI, a notable proportion (38.1%) of the food samples exhibited Class C β-lactamase ARGs detected by ARGpore2, whereas RGI detected only 5.4% of the samples (Figure 5E). However, RGI could identify additional carbapenem-resistant genes such as *bla^GES^, bla^GIM^, bla^IMI^, bla^IMP^, bla^KPC^, and bla^VIM^,* which were not detected by ARGpore2 (Figure 5E). Overall, RGI and ARGpore2 demonstrated varying profiles of ARG types and abundance across different food types.

### The influence of place of origin, retail venues and food processing methods on food microbiota

To assess the impact of various food attributes on the food microbiota, we identified distinct species associated with each food type based on factors such as place of origin (China, n=131; Europe, n=19; Asia-Pacific excluding China, n=73; North America, n=20; and South America, n=6), retail location (wet markets, n=38 or retail markets, n=222), and various food processing methods, including preservation (fresh, n=100; frozen, n=7; or chilled at 4°C, n=153), mincing (minced, n=46 or non-minced, n=134), and peeling (skin, n=46 or skinless, n=134) (Figure 6).

**Figure 6.**
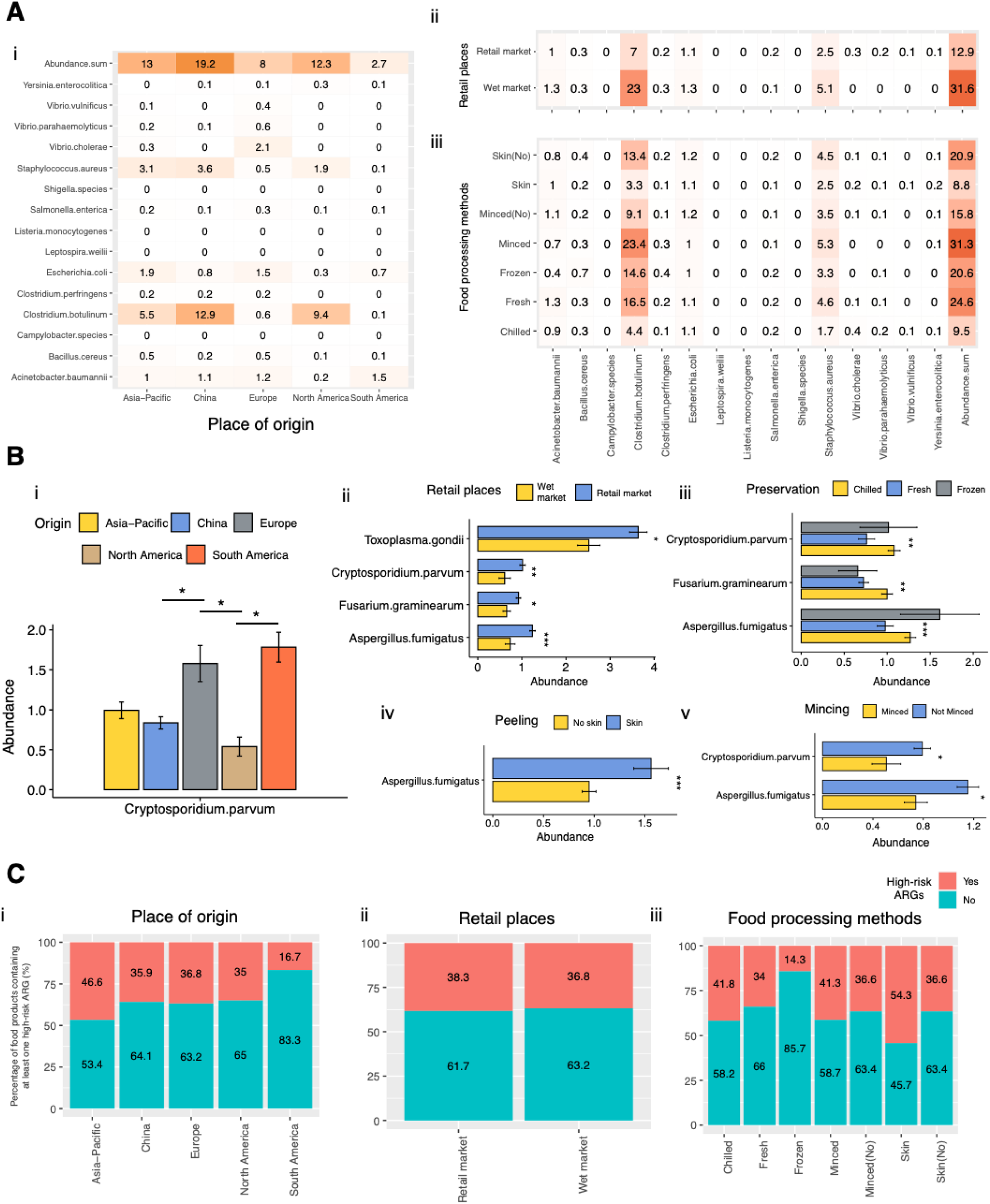
Examining the impact of origin, retail places and food processing methods on food microbiota and high-risk ARGs in food. (**A**) The mean abundance of 15 potentially pathogenic bacteria across different factors: (i) place of origin, (ii) retail places, and (iii) food processing methods such as preservation, peeling, and mincing. (**B**) The abundance of potential foodborne pathogens, encompassing protozoa (*Toxoplasma gondii* and *Cryptosporidium parvum*), fungi (*Fusarium verticillioides, Fusarium graminearum, Aspergillus fumigatus*, and *Aspergillus flavus*) within distinct food attributes: (i) place of origin, (ii) retail places, and food processing methods including (iii) preservation, (iv) peeling, and (v) mincing. Only significant results were shown. Significance levels are represented as *\*p < 0.05, **p < 0.01*, and *\*\*\*p <* 0.01 for all comparisons. (**C**) The percentage of food products containing at least one high-risk ARG across various food attributes, encompassing (i) place of origin, (ii) retail places, and (iii) processing methods.

We examined the abundance of 15 potentially pathogenic bacteria linked to various food attributes (Figure 6A). Notably, when considering their overall occurrence, China had the highest percentage at 19.2%, followed by the Asia-Pacific region at 13%, North America at 12.3%, Europe at 8%, and South America at 2.7% (Figure 6Ai). Foodborne pathogens such as *Clostridium botulinum* and *Staphylococcus aureus* were most commonly found in food imports from China, followed by North America and the Asia-Pacific (Figure 6Ai). In contrast, *Escherichia coli* had the highest prevalence in the Asia-Pacific and Europe, with a relative abundance of more than 1% (Figure 6Ai). Furthermore, our results indicated that a greater number of pathogenic bacteria were detected in wet markets compared to retail markets, particularly for *Clostridium botulinum* and *Staphylococcus aureus* (Figure 6Aii). Concerning food processing, higher levels of pathogenic bacteria were observed in fresh and frozen samples, accounting for 24.6% and 20.6%, respectively (Figure 6Aiii). Among these, *Clostridium botulinum, Staphylococcus aureus,* and *Acinetobacter baumannii* were the most abundant in fresh samples (Figure 6Aiii). As expected, both peeled and minced samples showed elevated levels of pathogenic bacteria, with *Clostridium botulinum* and *Staphylococcus aureus* being more prevalent in these samples (Figure 6Aiii).

In addition to investigating bacterial composition, we also examined how food attributes influenced the abundance of pathogenic protozoa (*Toxoplasma gondii* and *Cryptosporidium parvum*) and fungi (*Fusarium verticillioides, Fusarium graminearum, Aspergillus fumigatus,* and *Aspergillus flavus*). The results indicated a significantly higher abundance of *Cryptosporidium parvum* in food imported from Europe and South America, although no significant differences were observed for other species (Figure 6Bi). Concerning retail locations, a higher abundance of protozoa and fungi was found in retail markets compared to wet markets (Figure 6Bii). Additionally, food processing had a noticeable impact on the abundance of fungi and protozoa. Chilled samples exhibited significantly higher levels of pathogenic protozoa and fungi than fresh samples, although no significant differences were observed in frozen samples due to their small sample size (Figure 6Biii). Regardless of food handling, more *Aspergillus fumigatus* was observed in food with skin, while minced food exhibited higher levels of both *Aspergillus fumigatus* and *Cryptosporidium parvum* (Figure 6Biv and v).

### The influence of place of origin, retail place and food processing methods on high-risk ARG in food

The study also explored how different food attributes influenced the presence of high-risk ARGs in food (Figure 6C). The analysis revealed that food originating from the Asia-Pacific region exhibited the highest occurrence of high-risk ARGs, while food from South America had the lowest. Other regions displayed similar levels of high-risk ARGs (Figure 6Ci). Among these high-risk ARGs, the Asia-Pacific region had the highest abundance of genes in the carbapenem group (1.36), followed by China (0.99) and North America (0.80) (Table 1). Notably, only the Asia-Pacific region showed the presence of ARGs in the penam group (Table 1).

**Table 1.** Distribution of various drug classes linked to different food attributes, along with the average count of ARG reads in each sample.

|  | Carba-<br>penem | Cephalo-<br>sporin | Vanco-<br>mycin | Penam | High-risk<br>ARGs |
| --- | --- | --- | --- | --- | --- |
| **Place of origin** |  |  |  |  |  |
| Asia-Pacific | 1.36 | 0.33 | 0.03 | 0.01 | 1.73 |
| China | 0.99 | 0.11 | 0.00 | 0.00 | 1.11 |
| Europe | 0.68 | 0.16 | 0.05 | 0.00 | 0.89 |
| North America | 0.80 | 0.15 | 0.00 | 0.00 | 0.95 |
| South America | 0.33 | 0.00 | 0.00 | 0.00 | 0.33 |
| **Retail location** |  |  |  |  |  |
| Wet market | 1.08 | 0.18 | 0.00 | 0.00 | 1.26 |
| Retail market | 1.01 | 0.18 | 0.01 | 0.00 | 1.20 |
| **Food processing** |  |  |  |  |  |
| Chilled | 0.98 | 0.19 | 0.01 | 0.01 | 1.19 |
| Fresh | 1.11 | 0.17 | 0.01 | 0.00 | 1.29 |
| Frozen | 0.57 | 0.00 | 0.00 | 0.00 | 0.57 |
| Minced | 1.33 | 0.11 | 0.00 | 0.00 | 1.43 |
| Not Minced | 1.09 | 0.22 | 0.01 | 0 | 1.32 |
| No skin | 1.17 | 0.23 | 0.01 | 0.00 | 1.42 |
| Skin | 1.22 | 0.11 | 0.00 | 0.00 | 1.33 |

Interestingly, a similar pattern in high-risk ARGs was observed between food products sold in retail markets and those in wet markets (Figure 6Cii and Table 1). When considering food processing methods, we found that chilled samples had the highest proportion of high-risk ARGs at 41.8%, followed by fresh samples at 34.0%, and frozen samples at 14.3% (Figure 6Ciii). ARGs in the cephalosporin and vancomycin groups were detected in fresh and chilled foods exclusively (Table 1). Additionally, food with mincing and skin had a higher percentage of high-risk samples compared to non-minced and peeled samples (Figure 6Ciii). Intriguingly, ARGs in the carbapenem group exhibited higher abundance in samples with mincing and skin but lower abundance in the cephalosporin group compared to samples without mincing and skin (Figure 6Ciii and Table 1).

### Impact of sequencing depth on the microbiome taxonomic and AMR profiling

To examine how the sequencing depth affects microbiome profiling of food products, we randomly selected reads from each sample (>160 bp) and generated a new dataset with only 10% of the original sequencing depth, resulting in a reduced number of reads from 235130 ± 219214 to 23513 ± 21921 (Figure 7A, Supplementary Data 5). No bacterial read could be identified in three samples (Sample 137, 178, 260) after sequencing depth reduction which may be caused by its low bacterial count. We calculated the Sørensen–Dice coefficient between the original and reduced datasets using weighted and unweighted distances and found that the average coefficients for both distances were lower than 0.5 (Figure 7B, Supplementary Data 5). Partial Least Squares Discriminant Analysis (PLS-DA) was also performed to demonstrate potential differences between the two datasets (Figure 7C). The difference between the two datasets was less than 0.05 for both bacterial communities (difference score = 0.035), suggesting a small difference between them (Figure 7C).

**Figure 7.**
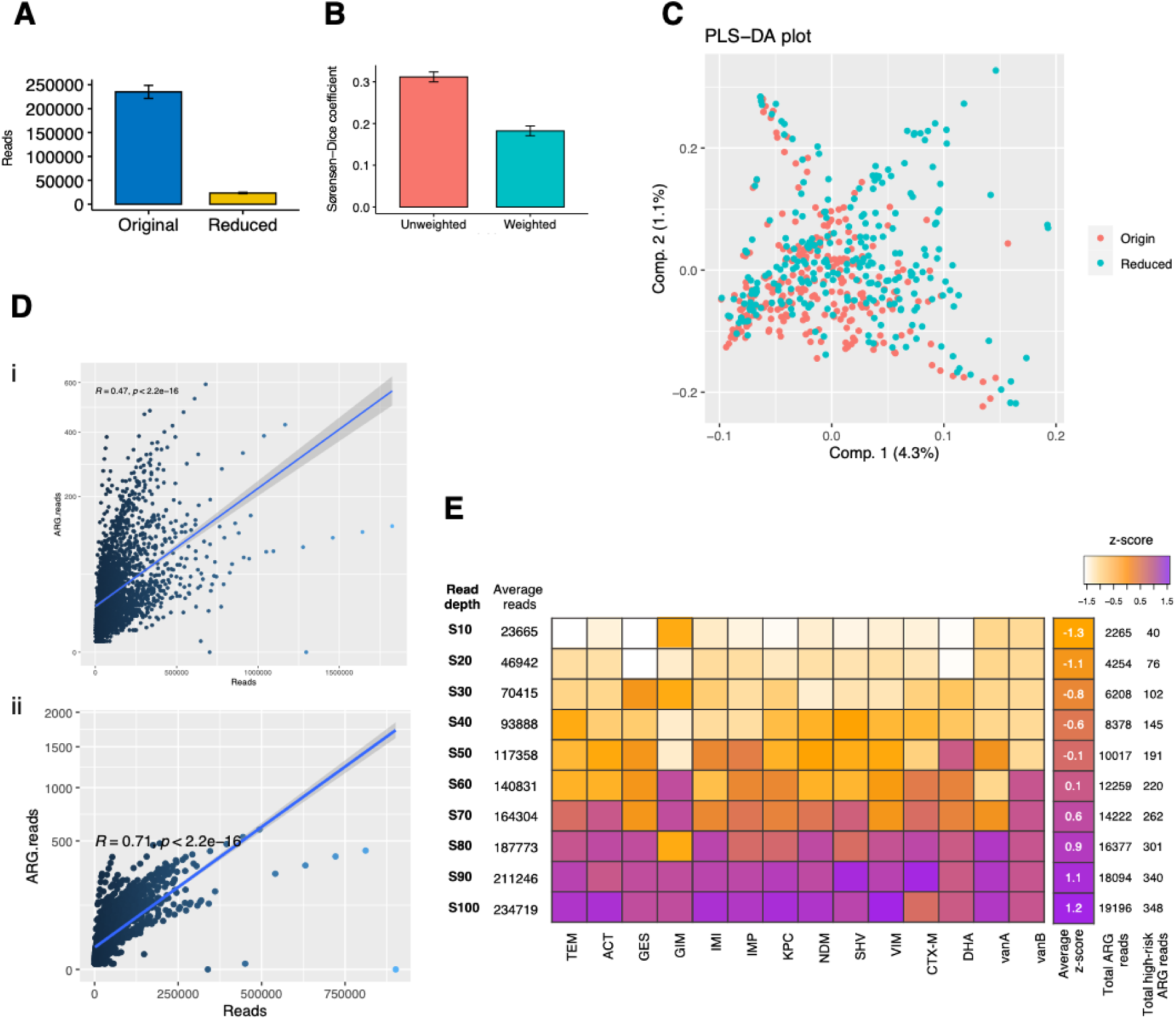
Impact of sequencing depth on the microbiome taxonomic and AMR profiling. (**A**) Read depth summary. The bar plot shows the total number of reads after random subsampling at 10% of the original dataset (Reduced) compared to the original dataset (Original). Data presented as mean ± SEM. (**B**) Sørensen–Dice coefficient between the original and reduced dataset using both weighted and unweighted distance metrics. (**C**) A PLS-DA plot of the bacterial communities (species of relative abundance of <10% were discarded) using the reduced and original dataset. (**D**) Correlation between (i) total reads and (ii) bacterial reads on the ARG reads. (**E**) Detection of high-risk ARGs among the randomly subsampled datasets. The high-risk ARG reads were analyzed using RGI. Multiple random subsampling are considered, including 90% (S90), 80% (S80), 70% (S70), 60% (S60), 50% (S50), 40% (S40), 30% (S30), 20% (S20), and 10% (S10) of the reads from the original dataset. The ARG profiles of these generated datasets are represented as relative abundances.

We subsequently investigated the effect of sequencing depth on ARG detection by using multiple randomly subsampled datasets. These datasets included different proportions of reads from the original dataset, ranging from 90% (S90) to 10% (S10). Figure 7D demonstrated notable correlations between the total number of reads and bacterial reads on the number of ARG reads (*p<0.05*). To determine the threshold where the effect size on ARG reads becomes less significant (effect size = 0.2), we identified threshold values of 703,034 and 338,973 for the total number of reads and bacterial reads, respectively. These values indicated the minimum number of reads required to ensure sufficient detection of ARGs. Furthermore, we examined the detection of high-risk ARGs with decreasing sequencing depth. We observed that when the proportion of original reads decreased to 60% (with an average total of 140,831 reads), the average z-scores showed a declining trend and dropped from 0.6 to 0.1 (Figure 7E). This indicates a deviation from the mean and a notable decrease in the detection of high-risk ARG reads.

### A comparison of AMR profiling and microbiome using Illumina and nanopore sequencing platforms in food samples

We analyzed thirty food samples that contained high-risk ARGs detected by nanopore sequencing and selected them for Illumina sequencing. The average size of the resulting nanopore (0.75 ± 0.52 G) and Illumina (8.61 ± 1.72 G) datasets are shown in Supplementary Data 6. After data processing, we observed a significant decrease in the average size of Illumina sequencing data (1.48 ± 0.53 G), while the size of nanopore sequencing data (0.72 ± 0.49 G) remained relatively unchanged (Figure 8A, Supplementary Data 6). Additionally, following sequence assembly in Illumina data, the average size was reduced to 0.46 ± 0.28 G. Cluster analysis of microbial compositions revealed that both platforms largely detected the same species. Data from the same samples generated by different platforms were clustered together, except for Sample021 (Supplementary Figure 1).

**Figure 8.**
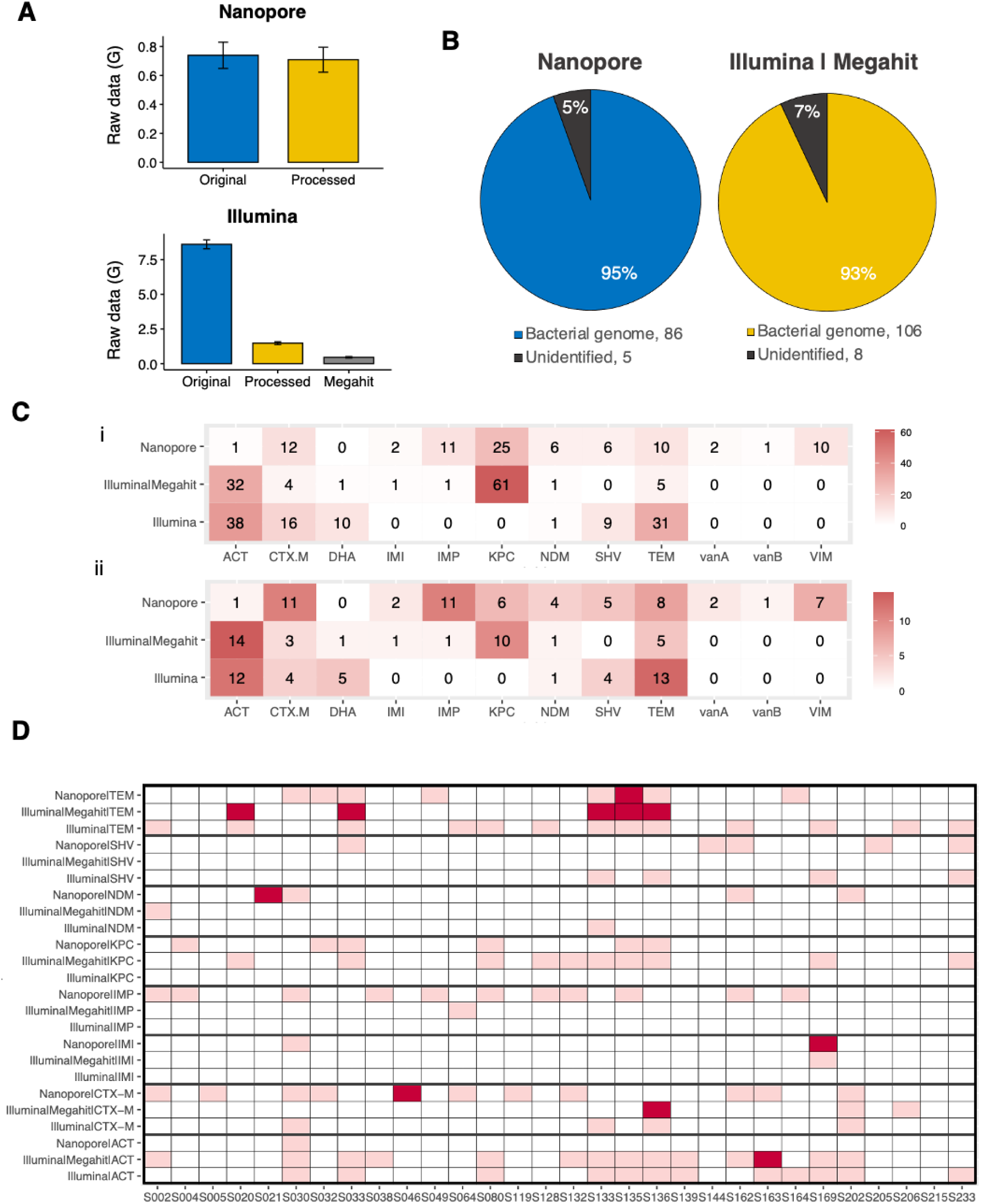
Performance comparison of Nanopore and Illumina metagenomic sequencing for AMR profiling in thirty food samples, both with and without sequence assembly by Megahit in Illumina data. (**A**) Raw data obtained from Illumina and Nanopore sequencing before and after processing. Data presented as mean ± SEM. (**B**) Percentages of high-risk ARG reads obtained from different platforms, mapped to bacterial genome. (**C**) Number of (i) ARG reads and (ii) samples classified as high-risk ARG in food products obtained from Illumina and Nanopore sequencing. (**D**) High-risk ARGs identified by Illumina and Nanopore sequencing in the 30 food samples. The food samples carrying ARGs and plasmid-mediated ARGs are in pink and dark red color, respectively.

Next, we employed RGI to identify high-risk ARGs from the Illumina sequencing data. The analysis of Illumina data with assembly revealed a higher number of high-risk ARGs with a total of 114 reads, whereas nanopore sequencing identified 91 reads of high-risk ARGs using the same database and ARG identifier (Figure 8B). Among the identified ARG reads, only 7% and 5% of the unidentified high-risk ARG reads were found in the Illumina and nanopore data, respectively (Figure 8B). Figure 8Ci presents the results of detecting specific high-risk ARGs using different sequencing methods: Illumina, Illumina with Megahit assembly, and Nanopore. The variability in the number of reads detected for different ARGs was observed in Illumina sequencing. Without assembly, Illumina data detected comparable reads (105 reads) with assembled Illumina data (106 reads) (Figure 8Ci). However, *bla^IMI^, bla^IMP,^ and bla^KPC^* genes were not detected in Illumina data without assembly. Interestingly, when Illumina was combined with Megahit assembly, the read counts for certain ARGs changed. The *bla^KPC^* gene showed an increase with 61 reads detected (Figure 8Ci). However, the *bla^SHV^* gene was not detected and *bla^CTX-M^, bla^DHA,^* and *bla^TEM^*read were reduced to > 4 folds in the assembled Illumina data (Figure 8Ci). Regarding *bla^VIM^*, *vanA*, and *vanB* genes, no reads were detected in both Illumina data with and without assembly (Figure 8Ci). In contrast, nanopore provided its own set of read counts for the ARGs. Both *vanA* and *vanB* genes were detected, but the *bla^DHA^* gene had no reads detected (Figure 8Ci). In terms of the number of samples in each high-risk ARG group, on average, 3 and 5 samples were identified in Illumina and Nanopore sequencing, respectively (Figure 8Cii). Nanopore sequencing detected more samples in all genes except *bla^ACT^, bla^KPC,^* and *bla^TEM^*(Figure 8Cii).

Figure 8D shows the high-risk ARGs detected by both sequencing platforms in the same food product (Supplementary Data 7 and 8). Only the ARGs that were identified by both platforms were displayed. Nanopore identified 59 ARG groups across all samples, whereas Illumina sequencing with and without assembly detected 36 and 39 ARG groups, respectively. Out of the 59 ARG groups identified by Nanopore, 12 of them (20.3%) were detected by both Nanopore and Illumina, regardless of assembly (Figure 8D). Intriguingly, different sequencing platforms identified distinct plasmids carrying ARGs, exhibiting variations in their sequences (Figure 8D). Furthermore, the accuracy of nanopore sequencing by aligning *bla^TEM^* genes generated by nanopore to those generated by Illumina was assessed (Supplementary Figure 2). For sequence lengths shorter than 300 bp, the accuracy was greater than 90%, while for sequence lengths longer than 800 bp, the accuracy was approximately 80% (Supplementary Figure 2).

## Discussion

Infections from foodborne pathogens, especially those involving AMR bacteria, continue to pose a significant threat to global health. Therefore, it has become crucial to explore and understand the microbial composition of different food products. Next-generation sequencing and nanopore sequencing technologies have advanced in recent years, enabling the study of food microbiomes using shotgun and metabarcoding sequencing methods (Gonçalves Dos Santos, Benito et al. 2017, Desdouits, Wacrenier et al. 2020, Li, Cao et al. 2020, Buytaers, Saltykova et al. 2021, Maguire, Kase et al. 2021). Our study provides the first comprehensive and cost-effective workflow that offers both screening and monitoring capabilities for food safety. We employed nanopore-based metagenomic sequencing to investigate the microbiome and AMR profiling of beef, chicken, pork, sashimi, and RTE-vegetables and successfully demonstrated its effectiveness and potential for enhancing food safety.

Our findings highlight that the microbiome profiles of meat and vegetables were distinct. Specifically, beef microbiome displayed a significant association with pork, while sashimi and chicken exhibited a strong correlation. The most dominant bacterial genus in beef was found to be *Clostridium*, while in pork, *Staphylococcus* and *Clostridium* were the most abundant genera. Interestingly, a previous study also indicated that traditional fermented pork from India was found to have a significant abundance of *Clostridium* species (De Mandal, Singh et al. 2018), which is also a natural contaminant of raw beef products (Novak and Yuan 2004). Previous studies have indicated that the presence of non-proteolytic *Clostridium* spores in meat could produce neurotoxins upon storage (Peck, Webb et al. 2020, Juneja, Purohit et al. 2021), highlighting the importance of monitoring *Clostridium sp.* in both pork and beef. Meanwhile, *Staphylococcus aureus*, a signature pathogenic bacterial species in pork, was identified as a potential food safety concern, with a previous culture-based study reporting its presence in 33.9% of pork products (Velasco, Vergara et al. 2018). In chicken, *Shewanella* was identified as the most dominant genus, with the species *Shewanella baltica* being the signature species, which can cause severe human infections such as bacteremia and pneumonia (Beaz Hidalgo, Agüeria et al. 2015). Similarly, in sashimi, *Shewanella* was the most dominant genus along with *Photobacterium*, with *Photobacterium damselae* being the most notable species. Although *Photobacterium damselae* is commonly found in seafood, it seldom causes infections in humans (Rivas, Lemos et al. 2013). However, during the processing of salmon, cross-contamination with *Pseudomonas* and *Shewanella* occurred, which raises concerns about the safety of sashimi (Møretrø and Langsrud 2017). In RTE-vegetables, the most prevalent bacteria were *Rahnella* and *Pantoea*, which are frequently found in packaged vegetable products with modified atmospheres (Franzetti, Musatti et al. 2015). It is important to note that these two species are opportunistic pathogens that can cause infections in salad vegetables (Hamilton-Miller and Shah 2001). Moreover, *Pseudomonas spp.* were among the signature species in RTE-vegetables, and previous research has also identified them as the most common bacteria in raw leafy salad vegetables (Ruiz-Roldán, Rojo-Bezares et al. 2021). Collectively, our findings align with previous studies, highlighting differences in food microbiomes. In addition, our study further pinpointed certain bacterial species that require careful surveillance when investigating particular types of food.

In terms of food safety, we conducted a thorough examination of the prevalence of foodborne pathogens and ARGs across various food types. Despite the previously mentioned prevalence of *Clostridium botulinum* and *Staphylococcus aureus* in different food types, we also observed that *Acinetobacter baumannii* was the dominant foodborne pathogen in RTE-vegetables. This finding is consistent with previous research indicating that vegetables can serve as a natural habitat for *A. baumannii*, a frequent cause of hospital infections (Berlau, Aucken et al. 1999). In addition, we detected the presence of *Yersinia enterocolitica*, a psychotropic foodborne pathogen, in meat samples. It is worth noting that meat can serve as a potential reservoir for *Y. enterocolitica*, thereby posing a significant public health concern (Fukushima 1985, Sirghani, Zeinali et al. 2018). In the case of sashimi, our findings concur with most existing literature, which highlights the predominance of *Vibrio species*. These bacteria have the potential to cause vibriosis and cholera (Maheshwari, Nelapati et al. 2011). Moving beyond bacteria, our investigation extended to other foodborne microorganisms, such as fungi and protozoa, within the food samples. Notably, our study unveiled a significant increase in the presence of *Toxoplasma gondii* in RTE-vegetables. This discovery aligns with previous research that emphasized the contamination of fresh vegetables with *T. gondii* oocysts (Marques, Sousa et al. 2020), further supporting our results.

In our antibiotic resistance genes (ARGs) analysis, we revealed a distinct pattern of ARGs across various food types. Of note, RTE-vegetables exhibited the highest abundance of high-risk ARGs, particularly those associated with carbapenem resistance genes. This pattern may be attributed to the excessive use of fertilizers and antibiotics in agriculture (Hudson, Frewer et al. 2017). Meanwhile, we successfully identified plasmids containing ARGs such as *bla^TEM^*, *bla^CTX-M^*, *bla^NDM,^*and *bla^IMI^* in diverse food samples. Specifically, plasmids carrying *bla^CTX-M^* genes were found exclusively in sashimi and RTE-vegetable samples. One of the sashimi samples harbored the *bla^CTX-M-12^*gene, conferring high-level resistance to cefotaxime, as investigated in a study (Bae, Lee et al. 2006). These transmissible resistance genes, present in the form of plasmids, can enable the transfer of drug-resistant characteristics from foodborne bacteria to the gut microbiota. Since the gut microbiota is often the primary source of bloodstream and urinary tract infections, it is essential to investigate ARGs from food samples, highlighting those mediated by plasmids. This allows us to understand and address the transmission dynamics of plasmid-mediated ARGs, mitigating the risks associated with AMR and its impact on human health. Taken together, our nanopore-based metagenomic approach could be applied as a food safety monitoring platform, using the abundance of pathogenic microbes and high-risk ARGs as indicators to determine the safety of food products. Although the total abundance of selected pathogenic microbes, including bacteria, fungi, and protozoa, was low in our findings, it is worth mentioning that some pathogens have a low infectious dose (Billington, Kingsbury et al. 2022). Furthermore, it is important to highlight that in RTE-foods, where thermal treatment is absent, the presence of bacterial or other microbial pathogens is more likely to cause infections. As such, the development of a standardized scoring system to assess food safety is necessary and warrants further investigation, particularly for RTE-food. Nevertheless, our approach has provided a foundational basis for food monitoring and is appropriate for surveillance purposes.

Furthermore, our study has provided a comprehensive understanding of how various food attributes, such as their places of origin, retail locations, storage conditions, and handling practices, impact microbial abundance. Notably, we observed that pathogenic species like *Clostridium botulinum* and *Staphylococcus aureus* were predominant in food samples imported from China. A review highlighted that these two species were among the pathogens responsible for a significant number of foodborne deaths in China (Wang, Duan et al. 2007). Concerning food processing methods, as expected, we noted higher levels of pathogenic bacteria in fresh, peeled, and minced food. This is attributed to the potential for contamination from sources such as unsanitary practices, leftover cuts and trims and the heat generated during mincing (Usda-Fsis 1999, Zhang, Singh et al. 2013, Kassem, Nasser et al. 2020). However, pathogenic bacterial abundance did not necessarily correlate with high-risk ARG abundance. Our findings revealed that chilled samples had a relatively high percentage of high-risk ARGs, even though the level of pathogenic bacteria was low. A previous study also indicated a higher prevalence of ESBL-producing Enterobacteriaceae in frozen foods compared to raw meat products, suggesting that storage procedures might be susceptible to cross-contamination during production (Ye, Wu et al. 2018). In addition, we highlighted the dominance of pathogenic protozoa like *Toxoplasma gondii* and *Cryptosporidium parvum*, as well as fungi including *Fusarium* and *Aspergillus spp.*, in specific food processing methods. In summary, our data provide valuable information for targeted monitoring of different food types subjected to varying food attributes when conducting metabarcoding sequencing or culture-based methods.

Prior studies have indicated that samples with high host DNA loads and low target abundance may require deeper sequencing, resulting in higher costs for metagenomic sequencing compared to metabarcoding sequencing (Feehery, Yigit et al. 2013, Billington, Kingsbury et al. 2022). Therefore, we investigated the effects of host DNA and sequencing depth on the results of metagenomic datasets. The similarity of the microbiome profile between the original and reduced datasets, which involved removing host DNA and reducing sequencing depth, remained largely unchanged overall. However, sequencing depth had a significant effect on ARG detection. These results are consistent with previous findings indicating that sequencing depth can substantially affect the AMR profiling of polymicrobial animal and environmental samples (Gweon, Shaw et al. 2019, Pereira-Marques, Hout et al. 2019). A minimum sequencing depth of one million Illumina reads is recommended for taxonomic classification, and 80 million Illumina reads are recommended for AMR identification (Gweon, Shaw et al. 2019). In the case of AMR profiling of wastewater using nanopore sequencing, approximately 120,000 metagenomic reads are recommended (Martin, Stebbins et al. 2021). Our findings suggest that reducing sequencing depth to approximately 703,034 (338,973 bacterial reads) and 140,831 nanopore reads could significantly impact the detection of ARGs and high-risk ARGs respectively. In summary, although reducing sequencing depth can lower the cost of food safety monitoring, it is important to use a higher sequencing depth for further AMR investigation of food samples to ensure accurate and reliable detection.

Moreover, we compared microbiome and AMR profiling in food samples using Illumina and nanopore sequencing platforms. Our results showed that both platforms could identify the same bacterial species, but the detection of ARGs was different. Nanopore sequencing was successful in detecting *vanA* and *vanB* genes, whereas Illumina sequencing specifically identified *bla^DHA^.* In line with these findings, a previous study indicated that nanopore-based metagenomics had limitations in detecting certain types of resistance genes, including *bla^VIM^, bla^KPC^, and bla^OXA^*genes (Viehweger, Marquet et al. 2021). Of note, the assembly process significantly influenced the detection of specific ARGs in Illumina. Previous studies have also highlighted the possibility of discrepancies between short-read, hybrid assemblies, and long-read assemblies in the patterns of ARGs (Brown, Keenum et al. 2021, Shuai, Itzhari et al. 2023). Apart from discrepancies in gene detection, the number of samples with high-risk genes detected by Illumina was lower. This discrepancy could be attributed to the reduction in the number of reads following the filtering of low-quality reads. In addition, it was suggested that Illumina may have higher accuracy than nanopore sequencing (Kerkhof 2021, Stevens, Creed et al. 2022), while we demonstrated an accuracy of over 90% for sequence lengths shorter than 300 bp and approximately 80% for sequence lengths longer than 800 bp in nanopore sequencing by aligning the TEM genes generated by nanopore to those generated by Illumina. Overall, we suggest that both platforms can be used for AMR profiling in food samples, but nanopore sequencing may be a better option.

In conclusion, our study has offered a promising workflow for food safety monitoring and emphasizes the need for careful surveillance of certain bacterial species when investigating specific types of food and food processing methods. We recommend using the abundance of pathogenic microbes and high-risk ARGs as indicators to determine the safety of food products, but a standardized risk assessment warrants further investigation. Furthermore, for efficient bacterial profiling in food samples, a lower sequencing depth may suffice for screening purposes, while a higher sequencing depth is necessary for AMR detection. Importantly, our findings suggest that nanopore sequencing may be a preferable choice for detecting ARGs when compared to Illumina sequencing. Taken together, this study provides a foundational basis for food monitoring and is appropriate for surveillance purposes. Future research can build on our approach to enhance food safety monitoring and establish a standard risk assessment to evaluate the potential risk to human health from foodborne AMR microorganisms.

## Materials and Methods

### Sampling and microbial extraction

A total of 260 food samples were collected from local markets (wet markets and retail markets) in Hong Kong, including 51 beef, 48 chicken, 81 pork, 23 sashimi, and 57 RTE-vegetable samples. Each food sample was cut and placed in a stomacher bag with 25 g of food content. Approximately 225 mL of sterile buffered peptone water was added to the bag and homogenized for 90 minutes at 230 rpm. After that, 50 mL of the homogenized contents were collected and transferred into a sterile 50 ml centrifuge tube. A bacterial pellet was obtained after centrifugation at 4,374 xg for 15 minutes. The pellet was then washed with sterile buffered peptone water and subjected to DNA extraction.

### DNA extraction

DNA was extracted from the bacterial suspension using the QIAamp BiOstic Bacteremia DNA Kit (Qiagen) according to the manufacturer’s protocol. In brief, DNA was released after bacterial cell lysis via mechanical homogenization. DNA was then purified by column-based methods after inhibitor removal procedures and collected with an elution buffer. The DNA content was then quantified using the Qubit 2.0 fluorometer (Thermo Fisher Scientific) with double-stranded DNA (dsDNA) HS assay kit (Thermo Fisher Scientific).

### Nanopore library preparation

Libraries were prepared using the Rapid Barcoding Kit (SQK-RBK110.96) from Oxford Nanopore Technologies (ONT) according to the manufacturer’s protocol and were quantified using the Qubit as described above. Ten barcoded libraries were then pooled with equal concentrations. After adapter ligation, sequencing was performed using the Flow Cell (R9.4.1) with the ONT GridION for 12h.

### Nanopore read data processing

Raw reads were trimmed by NanoFilt (De Coster, D’Hert et al. 2018). Reads with an average read quality score of <8.0 and of length below 1000 bp were discarded. The filtered clean reads were then classified by Kraken2/Bracken (Wood, Lu et al. 2019, Lu and Salzberg 2020) to determine the bacterial, archaeal, fungal, protozoal, and viral taxonomic compositions. Microbial species with a relative abundance of less than 1% were excluded from the analysis, except for the investigation of potential foodborne pathogens, which include 15 frequently studied foodborne pathogenic bacteria (*Acinetobacter baumannii, Bacillus cereus, Clostridium botulinum, Clostridium perfringens, Escherichia coli, Shigella species, Leptospira weilii, Salmonella species, Campylobacter species, Listeria monocytogenes, Staphylococcus aureus, Vibrio cholerae, Vibrio parahaemolyticus, Vibrio vulnificus, Yersinia enterocolitica)* (Prevention, Bottone 1999, Edwards 1999, Park 2002, Liu 2006, McIntyre, Bernard et al. 2008, Bağcıoğlu, Fricker et al. 2019, Elbehiry, Marzouk et al. 2021), protozoa (*Toxoplasma gondii* and *Cryptosporidium parvum*), and fungi (*Fusarium verticillioides, Fusarium graminearum, Aspergillus fumigatus*, and *Aspergillus flavus*) (Thrane 1999, Chang, Horn et al. 2014, Huerta-Cepas, Szklarczyk et al. 2019, Morris and Havelaar 2021). In parallel, the filtered reads were aligned to their corresponding host genome using the minimap2 tool (Li 2018). The reference genomes used for this alignment are listed in Table 2. Next, the unmapped reads were selected using SAMtools (Li, Handsaker et al. 2009), and a bacterial classification was performed on these reads.

**Table 2.** Reference genomes used for the alignment of nanopore reads.

| Host | Species name | Reference genome |
| --- | --- | --- |
| Beef | *Bos taurus* | GCF 002263795.1 |
| Bulb | *Allium sativum* | GCA 014155895.2 |
| Cabbage | *Brassica oleracea* | GCF 000695525.1 |
| Carrot | *Daucus carota sativus* | GCF 001625215.1 |
| Cherry tomato | *Solanum lycopersicum* | GCF 000188115.4 |
| Chicken | *Gallus gallus* | GCF 016699485.2 |
| Cucumber | *Cucumis sativus* | GCF 000004075.3 |
| Kale | *Brassica oleracea* | GCF 000695525.1 |
| Leek | *Allium cepa* | GCA 905187595.1 |
| Lettuce | *Lactuca sativa* | GCF 002870075.3 |
| Onion | *Allium cepa* | GCA 905187595.1 |
| Pork | *Sus scrofa* | GCF 000003025.6 |
| Salmon | *Salmo salar* | GCF 905237065.1 |
| Scallop | *Pectinidae* | GCF 902652985.1 |
| Shrimp | *Pandalus platyceros* | GCA 005815305.1 |
| Spinach | *Spinacia oleracea* | GCA 000510995.2 |
| Sweet pepper | *Capsicum annuum* | GCF 002878395.1 |
| Tuna | *Thunnus albacares* | GCF 914725855.1 |
| Yellowtail | *Seriola quinqueradiata* | GCA 002217815.1 |

### Antimicrobial resistance profiling using nanopore reads

For the identification of ARGs in the food metagenomic data, a selection process was applied to the sequenced reads. Specifically, reads were chosen based on two criteria: (1) an average read quality score above 8.0 and a minimum length of 160 bp for the Resistance Gene Identifier (RGI) analysis, and (2) a minimum length of 1000 bp for the ARGpore2 analysis (Alcock, Raphenya et al. 2020, Cheng 2022). CARD was run using the default settings and CARD’s Resistance Gene Identifier (RGI) 4.2.2 with the CARD 3.0.1 database.

Only “Perfect” and “Strict” AMR hits were included as determined by curated similarity cut-offs. ARG with a minimum percentage length of reference sequence of 30% and a minimum sequence identity of 80% were selected (Whittle, Yonkus et al. 2022). AMR profiles analyzed by ARGpore2 were generated by default parameters and the database. Reads that were defined as high-risk ARGs, including key β-lactamase genes, *bla^TEM^*, *bla^ACT^*, *bla^GES^*, *bla^GIM^*, *bla^IMI^*, *bla^IMP^*, *bla^KPC^*, *bla^NDM^*, *bla^SHV^*, *bla^VIM^*, *bla^CTX-M^*, *bla^DHA^*, glycopeptide genes (*vanA* and *vanB*), as well as methicillin resistance gene (*mecA*), were isolated (Zhang, Gaston et al. 2021). Only sequences that were mapped to a bacterial genome using Nucleotide BLAST 2.13.0+ (Camacho, Coulouris et al. 2009) were used for further investigation. ARGs classified as plasmids by Nucleotide BLAST 2.13.0+ were also extracted (Camacho, Coulouris et al. 2009).

### Illumina sequencing

Thirty DNA samples, each containing at least one high-risk ARG, were subjected to library preparation, quality assessment and Illumina sequencing. In brief, the library was prepared using the NEBNext Ultra RNA Library Prep Kit for Illumina (Cat No. 7530) according to the manufacturer’s protocol. The sample quality was then assessed using NanoDrop, agarose gel electrophoresis and the Agilent 2100 BioAnalyzer (Agilent Technologies, Palo Alto, CA, USA). For sequencing, Illumina Novaseq 6000 (Illumina, San Diego, CA, USA) with 150 bp paired-end was performed.

### Illumina read data processing

Raw reads were processed using TrimGalore-0.6.6 (a wrapper script for Cutadapt and FastQC) for adapter and quality trimming (F. 2015). After trimming, reads longer than 160 bp were subjected to taxonomic classification using Kraken2/Bracken (Wood, Lu et al. 2019, Lu and Salzberg 2020) to determine the bacterial composition of the sample. SeqKit was utilized for read statistics and read selection procedures (Shen, Le et al. 2016). Microbial species with a relative abundance of less than 1% were excluded, except for the investigation of potential foodborne pathogens. For ARG detection, only reads that were longer than 160 bp were considered using the RGI with the BWT (Burrows-Wheeler Transform) mode (Alcock, Raphenya et al. 2020). For the output, mapped reads with MAPQ > 70, and coverage > 30 % of the reference length were selected for downstream analysis (Shuai, Itzhari et al. 2023). Meanwhile, the trimmed reads were assembled using Megahit, a fast and memory-efficient de novo assembler (Li, Liu et al. 2015). The assembled reads were subjected to ARG detection using RGI, following a similar approach as the processing of nanopore reads.

### Principal component analysis, Partial Least Squares Discriminant Analysis, correlation analysis and hierarchical clustering

R-based computational tools in R-studio were used to generate the graphs (McMurdie and Holmes 2013, McMurdie and Holmes 2014, McMurdie and Holmes 2015, Callahan, Sankaran et al. 2016). Principal component analysis (PCA) was performed with the R prcomp function from the stats package on a Bray-Curtis dissimilarity matrix calculated with the R vegdist function from the vegan package (Dixon 2003, Holland 2008). Two principal components were retained for plotting. The resulting PCA plot displayed the distribution of microbiomes of the average bacterial abundance from five types of food samples in the dataset, with bacterial species with an abundance greater than 1% included. Using the same dataset, a correlation matrix heatmap was generated to further explore the relationships between the microbiomes of the five food types using the ggplot2 package in R. The heatmap visually displays the correlation coefficients between pairs of food products. The hierarchical clustering heatmap was generated using the R package “pheatmap” (Kolde and Kolde 2018). For Partial Least Squares Discriminant Analysis (PLS-DA), the R pls function package in R was used to analyze the discrepancy between the original and reduced datasets (Mevik and Wehrens 2015). A scatter plot of the PLS-DA scores was generated using ggplot2. To calculate the difference between the two datasets in the plot, the mean scores for each PLS-DA component in each group were calculated.

### Identification of signature species

The multilevel pattern analysis was performed using the R package “indicspecies” to identify bacterial species that were significantly associated with different sample groups (De Caceres, Jansen et al. 2016). The association function “r.g” was applied with a significance level of 0.05. The “r.g” function specifically calculates the correlation coefficient (r) between each species and each site group, and tests for statistical significance using a permutation test. Species with a test statistic value greater than 0.25, which was calculated for the association between groups, were classified as signature species (Bergerot, Fontaine et al. 2011).

### Concordance of the microbiomes between samples

Sørensen–Dice coefficient was used to measure the similarity between microbiota among the samples, based on the presence and absence of species (unweighted) and the abundance of species (weighted). For each sample pair, the Sørensen index was calculated using the function “vegdist” with the “bray” method in the R package (Dixon 2003). The Sørensen index is a distance metric that ranges between 0 and 1, where a value of 0 indicates complete similarity between the two sets of samples, and a value of 1 indicates no similarity.

## Statistical analysis

To assess the effects of sequencing depth on ARG detection, all statistical comparisons, *\*p < 0.05 **: p < 0.01 ***: p < 0.001* were used as the significance level for all comparisons (Kruskal-Wallis test).

## Data availability

Sequence data were archived in the National Center for Biotechnology Information (NCBI) Short Read Archive (SRA) (PRJNA991711).

## Conflict of Interest

The authors declare that the research was conducted in the absence of any commercial or financial relationships that could be construed as a potential conflict of interest.

## Author Contributions

AL and GS conceived the original concept and designed the study. AL, EW, IN, IW, RS, DC, VW, ZZ, SF, SW, WT, and HL performed the experiments. AL analyzed the data and wrote the original paper. GS and AL amended and revised the writing. All authors read and approved the final version of the manuscript.

## Funding

This work was supported by Health and Medical Research Fund (#23220402).

---

**Supplementary Figure 1.**
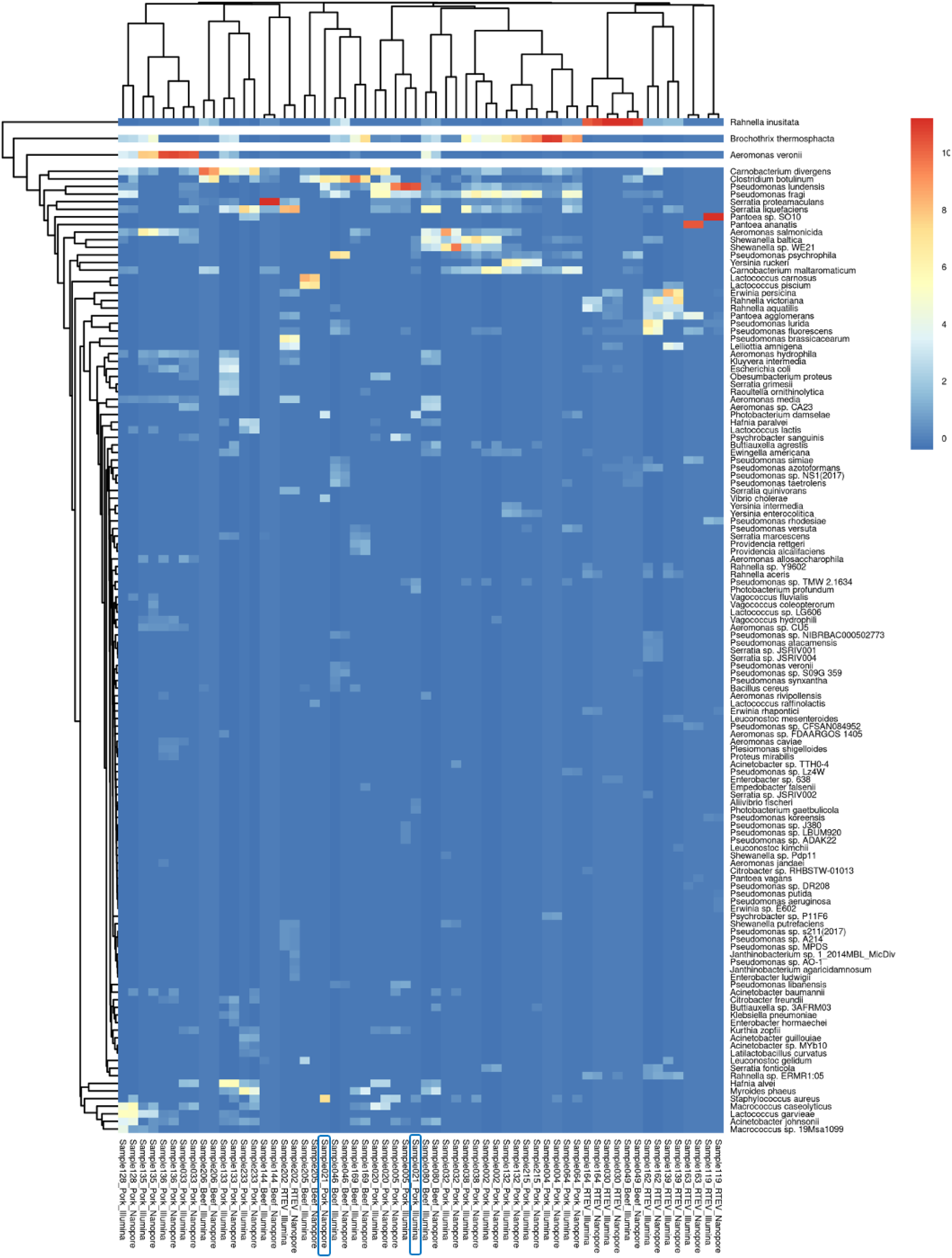
Microbial compositions of different food products based on the sequencing data. The blue frame highlights the dissimilarity observed in the microbiome between the same food products when different sequencing platforms are employed.

**Supplementary Figure 2.**
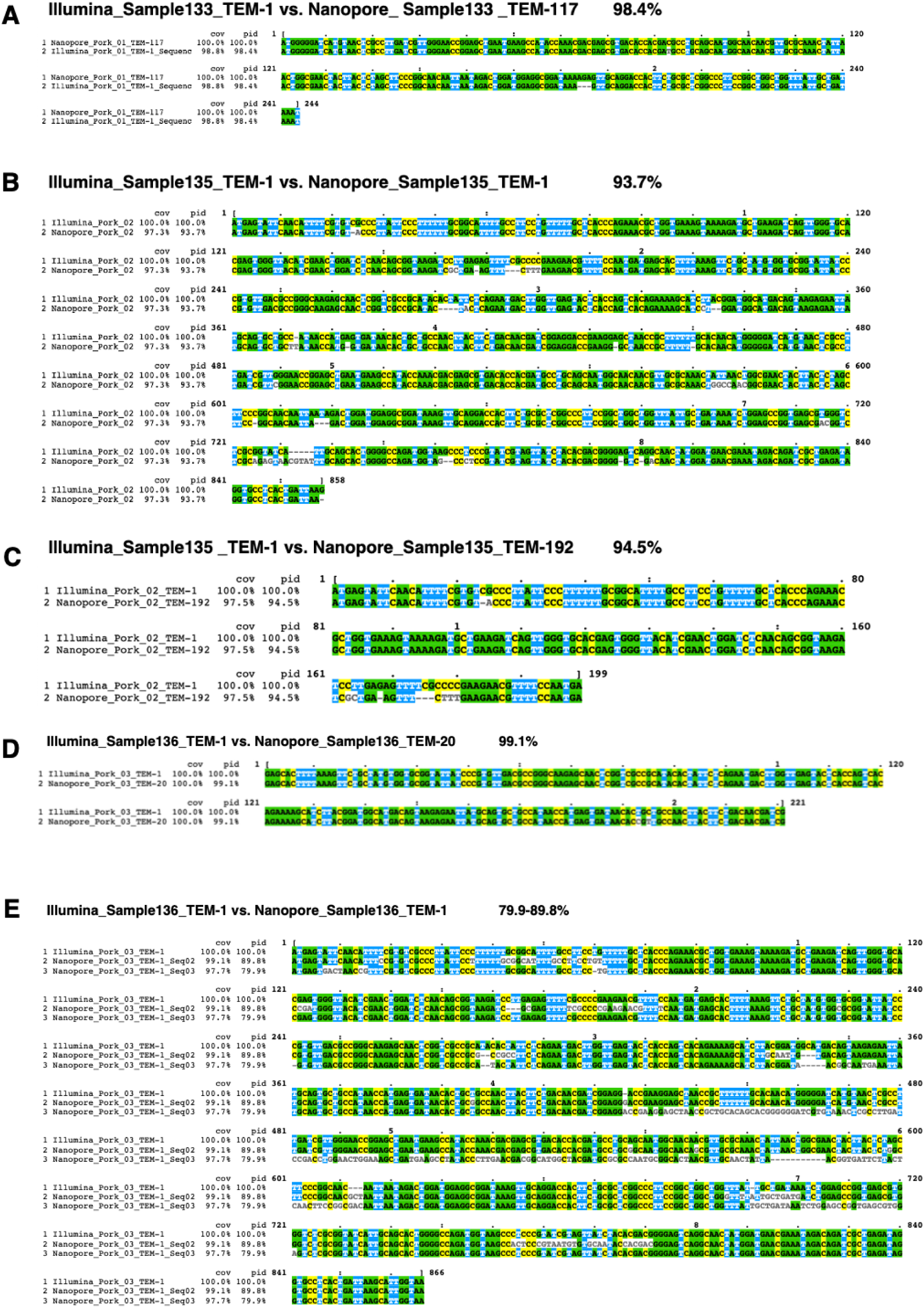
Sequence alignment of TEM genes. (**A**) TEM-117, (**B**) TEM-1, (**C**) TEM-192, (**D**) TEM-20, (**E**) TEM-1, generated by Nanopore and Illumina metagenomic sequencing.

## Notes

### Competing Interest Statement

The authors have declared no competing interest.

